# Changes in ADAR RNA Editing Patterns in CMV and ZIKV Congenital Infections

**DOI:** 10.1101/2023.06.16.545385

**Authors:** Benjamin Wales-McGrath, Heather Mercer, Helen Piontkivska

## Abstract

**Background:** RNA editing is a process that increases transcriptome diversity, often through Adenosine Deaminases Acting on RNA (ADARs) that catalyze the deamination of adenosine to inosine. ADAR editing plays an important role in regulating brain function and immune activation, and is dynamically regulated during brain development. Additionally, the ADAR1 p150 isoform is induced by interferons in viral infection and plays a role in antiviral immune response. However, the question of how virus-induced ADAR expression affects host transcriptome editing remains largely unanswered. This question is particularly relevant in the context of congenital infections, given the dynamic regulation of ADAR editing during brain development, the importance of this editing for brain function, and subsequent neurological symptoms of such infections, including microcephaly, sensory issues, and other neurodevelopmental abnormalities. Here, we begin to address this question, examining ADAR expression in publicly available datasets of congenital infections of human cytomegalovirus (HCMV) microarray expression data, as well as mouse cytomegalovirus (MCMV) and mouse/ human induced pluripotent neuroprogenitor stem cell (hiNPC) Zika virus (ZIKV) RNA-seq data.

**Results:** We found that in all three datasets, ADAR1 was overexpressed in infected samples compared to uninfected samples. In the RNA-seq datasets, editing rates were also analyzed. In all mouse infections cases, the number of editing sites was significantly increased in infected samples, albeit this was not the case for hiNPC ZIKV samples. Mouse ZIKV samples showed altered editing of well-established protein-recoding sites such as Gria3, Grik5, and Nova1, as well as editing sites that may impact miRNA binding.

**Conclusions:** Our findings provide evidence for changes in ADAR expression and subsequent dysregulation of ADAR editing of host transcriptomes in congenital infections. These changes in editing patterns of key neural genes have potential significance in the development of neurological symptoms, thus contributing to neurodevelopmental abnormalities. Further experiments should be performed to explore the full range of editing changes that occur in different congenital infections, and to confirm the specific functional consequences of these editing changes.

## Background

Congenital infections of the unborn fetus or newborn infant are a widespread issue caused by pathogens such as the TORCH pathogens and other emerging pathogens such as Zika virus (ZIKV) (Boucoiran et al., 2020; Coyne and Lazear, 2016). These frequently disrupt fetal development, particularly in the brain, resulting in a wide range of symptoms, including damage to sensory organs, birth defects such as microcephaly, or learning and behavioral disorders (Boucoiran et al., 2020; Canetta et al., 2016). Pathogens often target neural stem cells (NSCs) and neural progenitor cells (NPCs), inducing apoptosis, curtailing proliferation, and impairing the differentiation of these cells (Boucoiran et al., 2020; Kamte et al., 2021). Interestingly, these effects are often caused, not only by the direct influence of infection, but as the indirect result of the immune response, as cytokines and chemokines have been shown to have significant effects on NSC/NPC proliferation, survival, and differentiation (Boucoiran et al., 2020; Chandwani et al., 2019; Kamte et al., 2021). However, the mechanisms by which infections cause these effects remain to be fully elucidated, and can be highly dependent on the stage of pregnancy where infection occurs (Boucoiran et al., 2020; Kamte et al., 2021). Here we will focus on the effects of two of the most common congenital infections, cytomegalovirus (CMV) and ZIKV.

### Cytomegalovirus

Cytomegalovirus (CMV), a dsDNA virus of the family Herpesviridae, is the most common cause of congenital infection, with primary infection occurring in 0.7-4% of pregnancies (Nigro, 2009). The most common symptoms in congenitally infected children include sensorineural hearing loss (SNHL), microcephaly, cerebral calcifications, white matter lesions, and other central nervous system (CNS) abnormalities (Guerra et al., 2008; Korndewal et al., 2017; Pesch et al., 2021; Picone et al., 2014; Vande Walle et al., 2021). This can then lead to motor, cognitive, and speech-language developmental delays, though symptoms can depend on the stage of pregnancy during which infection occurs (Korndewal et al., 2017; Pass et al., 2006; Pesch et al., 2021). Neurological symptoms of congenital CMV infection result from the virus’s tropism for NSCs and radial cells (Teissier et al., 2014). Once in the CNS, CMV can cause damage by inducing apoptosis, reducing proliferation, and impairing differentiation in NSCs and NPCs (Kamte et al., 2021; Zhang and Fang, 2019). CMV has been reported to cause transcriptional changes that dysregulate neurodevelopment (Liu et al, 2017; Rolland et al., 2016), but the induction of an inappropriate immune response by CMV has also been implicated in the pathogenesis of neurodevelopmental symptoms (Cheeran et al., 2009). While CMV infection has been shown to induce peripheral leukocyte infiltration, microglial activation, and increased chemokine expression (Cloarec et al., 2016; Seleme et al., 2017), attenuation of inflammation with TNF-α (TNFA)-neutralizing antibodies was shown to reduce neurodevelopmental abnormalities with little effect on viral replication (Seleme et al., 2017). However, the exact mechanisms behind these effects are unknown.

### Zika Virus

ZIKV is a positive-sense ssRNA virus of the Flaviviridae family (Fields et al., 2013; Baud, 2017; Piontkivska et al., 2017). ZIKV is primarily transmitted by *Aedes* mosquitoes (Baud et al., 2017; Faye et al., 2014) and distributed throughout sub-Saharan Africa and Southeast Asia. In 2015-2016 it caused a major public health emergency in Brazil and other countries worldwide (Baud et al., 2017). Importantly, transmission from mother to fetus can also occur, resulting in congenital infection, including tropism for fetal brain tissue (Baud et al., 2017; Bhatnagar et al., 2017; Brasil et al., 2016; Hoen et al., 2018; de Noronha et al., 2016 ; Reynolds et al., 2017; Rosenfeld et al., 2017). These infections are associated with a number of adverse neurological consequences for the fetus, termed congenital Zika syndrome (CZS), which develops in up to 15% of pregnancies (Hoen et al., 2018; Moore et al., 2017; Reynolds et al., 2017). One of the most severe features of CZS is microcephaly (a condition characterized by significantly reduced head size), as well as other abnormalities in cranial morphology (Brasil et al., 2016; Landry et al., 2017; Moore et al., 2017; Rasmussen et al., 2016). In addition to this physical damage, CZS may be accompanied by more long-term neurodevelopmental consequences, although data on long term developmental outcomes from children with CZS is relatively sparse (Moore et al., 2017; Nielsen-Saines et al., 2019; Pecanha, et al., 2020; Stringer, et al. 2021).

ZIKV can infect NSCs and NPCs, disrupting their survival, proliferation, and differentiation, as well as neuronal migration, thus impairing brain development and neurogenesis (Chang et al., 2020; McGrath et al., 2017; Rosenfeld et al., 2017; Tang et al., 2016). Interestingly, apoptosis is induced not just in cells infected by ZIKV, but in uninfected cells in the surrounding area, indicating the significance of cytotoxic factors not directly resulting from viral infection (Rosenfeld et al., 2017). While the mechanisms by which ZIKV causes these effects are not well understood, it has been shown to dysregulate gene expression in pathways related to brain development and cell death, and interestingly, immune response and microglial activation, including ADAR (Chang et al., 2019). Further evidence suggests that ZIKV induces a strong type 1 interferon (IFN) response and may play a role in ZIKV pathogenesis (Lima et al., 2019).

### What are ADARs and What do they Do?

ADAR enzymes are the primary cause of RNA editing, catalyzing the deamination of adenosine to inosine in dsRNA. The ADAR family contains three gene loci in mammals: ADAR1, ADAR2, and ADAR3 (Piontkivska et al., 2021; Savva et al., 2012; Jin et al., 2009). Of these, only ADAR1 and 2 are known to engage in editing, as ADAR3 is believed to be catalytically inactive and can inhibit editing by ADAR1 and 2 (Nishikura et al., 2016; Tan et al., 2017; Oakes et al., 2017). ADAR1 has two isoforms (p110 and p150), which are transcribed from unique promoters, the latter of which is IFN-inducible due to an interferon-sensitive response element (ISRE) (George and Samuel, 1999; Samuel, 2019). In addition, ADAR1p110 and ADAR2 are primarily localized to the nucleus, while ADAR1 p150 can be exported to the cytoplasm (Nishikura et al., 2016; Savva et al., 2012). Cellular localization may contribute to differences between RNA editing targets of ADAR1 p150 and ADAR1 p110/ADAR2, particularly in the context of editing of cytoplasmic dsRNA by ADAR1 p150 upregulated as part of the innate immune response (Nishikura et al., 2016; Yanai et al., 2020; Cruz et al., 2020).

Editing by these enzymes can take the form of highly selective site-specific editing or less selective hyper-editing of multiple bases in a single transcript. The former is more likely to occur in shorter duplexes with imperfect base pairing, while the latter occurs more frequently in longer double-stranded transcripts (Savva et al., 2012). Most of these editing sites occur in non coding transcripts, while a small, but disproportionately important minority of editing sites occur in protein coding RNA sequences (Nishikura, 2016; Samuel, 2019). Specifically, noncoding editing most commonly takes place in transcripts of repetitive sequences, primarily *Alu* sequences in humans (Samuel, 2019), as well as in intergenic regions, introns, and 3’ UTRs (Chen, 2013). The nucleus contains a higher proportion of total editing sites than cytosol, specifically for editing sites in introns or intergenic regions, while 3’ UTR editing sites are disproportionately found in the cytosol (Chen, 2013). Editing also varies by tissue and by brain region, possibly due to differences in ADAR expression, or differences in expression of other trans-acting regulatory factors (Tan et al., 2017).

This editing can affect protein coding, as well as splicing, miRNA binding, and other non-coding RNAs. While the number of protein recoding editing sites is relatively small, many of these sites are highly conserved and have physiologically significant effects (Nishikura, 2016; Hood and Emeson 2012; Peng et al., 2012). This is particularly significant with editing of neurological genes such as GRIA2 (encoding the glutamate ionotropic receptor AMPA type subunit 2 and edited to produce a Q/R substitution) and HTR2C (encoding the G protein-coupled serotonin receptor 5-HT2CR and edited at 5 sites) (Nishikura, 2016; Zhai et al., 2022). Dysregulation of this editing can cause neuron cell death in the case of GRIA2 (Bass, 2002; Higuchi et al., 2000), or more nuanced effects such as altered sensitivity to neurotransmitter signaling in the case of HTR2C (Bass, 2000; Nishikura, 2016; Slotkin and Nishikura, 2013). Neural ADAR editing has been linked to neurological diseases, including amyotrophic lateral sclerosis (ALS) and Alzheimer’s disease (Lorenzini et al., 2018; Slotkin and Nishikura, 2013), as well as Prader-Willi Syndrome (Slotkin and Nishikura, 2013; Kishore and Stamm, 2006). Editing changes have also been implicated in psychiatric disorders, such as anxiety, depression, and schizophrenia (Lorenzini et al., 2018; Slotkin and Nishikura, 2013; Maas et al., 2006; Breen et al., 2019).

Aside from protein recoding, ADAR editing can affect other aspects of RNA regulation, such as alternative splicing, through changing splice site sequences, editing of splicing regulatory elements (SREs), or altering RNA secondary structure near splicing junctions (Kapoor et al., 2020; Savva et al., 2012; Solomon et al., 2013; Tang et al., 2020). ADAR may also have editing-independent effects on splicing, for example, through competing with splicing factors for RNA binding or by recruiting splicing factors (Tang et al., 2020). Additionally, ADAR editing of miRNAs or miRNA binding sites in 3’ UTRs can influence miRNA targeting, through changes in processing of primary miRNAs (pri-miRNAs) to mature miRNAs, including through both editing of pri-miRNAs and through interaction of ADAR1 with Dicer, an enzyme responsible for precursor miRNAs (pre-miRNA) cleavage (Nishikura, 2016; Ota et al., 2013; Yoshida et al., 2021; Wang Q. et al., 2013). Because of this, changes in ADAR expression can lead to changes in the abundance of miRNAs and their targets (Vesely et al., 2012). Importantly, gene regulation by ADAR enzymes is subject to dynamic and nuanced regulation during human and mouse brain development, and in disease states (Deffit and Hundley 2016; Heraud-Farlow et al., 2020; Breen et al., 2019). Overall, regulation of ADAR editing in the brain is complex, with nuanced changes over development and across brain regions, and with a dynamic interplay between different ADAR enzymes which remains to be fully understood (Deffit and Hundley 2016; Heraud-Farlow et al., 2020; Breen et al., 2019). Additionally, this regulation can be particularly difficult to untangle once additional complicating factors such as immune responses are involved, as seen in congenital viral infection or maternal immune activation.

Aside from its function in gene regulation, ADAR enzymes (primarily ADAR1 p150 isoform) play a dual role in the antiviral innate immune response. As noted previously, expression of ADAR1 p150 is induced by signaling from type 1 IFNs, a class of cytokine secreted in response to viral infection (Ivashkiv and Donlin, 2013; MacMicking 2012). During the antiviral immune response, editing has complex pro- or anti-viral effects, dependent on host/virus-specific factors or even changing over the course of infection. The variable effects of this editing can disrupt the function of viral proteins, while suppressing interferon responses or potentially giving rise to variants that enable immune escape (Piontkivska et al., 2021; Pfaller et al., 2021).

**Evidence of ADAR1 p150 activation by IFNs leads to the question: what impact does the induction of ADAR1 p150 expression in response to viral infection have on normal host transcriptome editing?** This question is particularly significant in the brain, and in the context of congenital infection, given the importance of proper regulation of ADAR editing for brain development. However, a handful of prior studies showed conflicting results as to the effect of infection on ADAR editing rates (reviewed in Piontkivska et al., 2021). While infection of primary human neural stem cells with Zika virus has been shown to cause increased ADAR editing (including in GRIA3 recoding site) (Piontkivska et al., 2019), infection of neonatal mice by a neurotropic strain of reovirus (ReoV) showed that, despite a strong induction of ADAR1 p150, editing changes were limited to a small number of sites (Hood et al., 2014).

In addition, maternal immune activation (MIA) using poly(I:C) in mice to mimic the immune response to viral infection, has been used to investigate the congenital effects on ADAR editing; the results showed increased global editing, including in protein recoding sites for significant neurological genes. Although these editing increases were temporary, as sequencing later in life showed little difference between polyI:C and controls, the MIA-induced behavioral abnormalities persisted (Tsivion-Visbord et al., 2020). These results suggest that even transient changes in editing patterns can have far-reaching functional consequences (Piontkivska et al., 2019), underscoring the need to understand how the most common sources of congenital infection affect ADAR editing, and how editing dysregulation contributes to neurodevelopmental symptoms later in life.

Here, we explore whether congenital viral infection is associated with changes in ADAR editing of key host genes, and whether genes with editing changes can be linked to important neurodevelopmental functions, by examining publicly available microarray data of congenital human cytomegalovirus (HCMV) infection and RNA-seq data of congenital mouse cytomegalovirus (MCMV) infection and Zika virus (ZIKV) infection in mice to assess the effect of congenital infections on ADAR expression and editing. Our results show increased ADAR1/2 expression and increased A-to-I editing associated with infection, including changes in editing in genes relevant to neurodevelopment, providing support for the hypothesis that dysregulation of ADAR editing contributes to neurodevelopmental abnormalities caused by congenital infection.

## Results

To examine the effects of congenital infections on ADAR editing in the developing brain, we examined transcriptome datasets from humans or model organisms with congenital infections. Using the NCBI BioProject database, we identified 5 such relevant datasets with humans and mice infected with ZIKV and CMV, as described in Table 1.

**Table 1.** List of BioProject datasets of viral infections used in this study, and their characteristics.

| BioProject Accession | Virus | Time of Infection | Organism | Virus Delivery | Tissue | ADAR1 Expression | ADAR2 Expression | ADAR3 Expression | Editing Site Number | Editing Sites with Increased Editing | Editing Sites with Decreased Editing |
| --- | --- | --- | --- | --- | --- | --- | --- | --- | --- | --- | --- |
| PRJNA422858 | HCMV<sup>#</sup> | Infants at time of diagnosis | Human | Natural | Blood | Up (p<sub>adj</sub> = 7.31E-07) | Up (p<sub>adj</sub> = 7.59E-05 and 4.00E-03) | No Change | - | - | - |
| PRJEB38849 | MCMV | Newborn mice, analyzed after 8 days infection | Mouse | inoculated i.p. with 400 PFU | Microglia | Up (p = 0.007949) | No Change | No Change | Up (p = 0.01129) | 21 | 0 |
| PRJNA487357 | ZIKV | E10.5/E14.5 | Mouse | unspecified | Brain | Up (p = 0.0005574) | No Change | No Change | Up (p = 0.01877) | 12 | 0 |
| PRJNA358758 | ZIKV | E15.5/P3 | Mouse | 6.5 × 10<sup>5</sup> PFU injected into one side of the lateral ventricle (LV) | Brain | Up (p = 0.03035) | No Change | Down (p = 0.01266) | Up (p = 0.0007137) | 137 | 11 |
| PRJNA551246 | ZIKV | Analysis after<br>48 hours<br>infection | Human | Inoculated for<br>1hr with virus<br>at MOI 1 | hiNPC | Higher in<br>PE243V<br>(p_adj =<br>1.81e-05)<br>and FSS1302<br>(p_adj =<br>3.00e-06)<br>than control;<br>higher in<br>FSS1302<br>than PE243V<br>(p_adj =<br>7.22e-03) | Higher in<br>FSS1302<br>than controls<br>(p =<br>0.0490919) | No Change | No Change | 3 in<br>FSS1302; 1<br>in PE243V;<br>1 in both | 2 in<br>FSS1302; 1<br>in PE243V;<br>1 higher in<br>PE243V<br>than<br>FSS1302 |

Characteristics of BioProject datasets of infections of Zika virus, ZIKV, and human and mouse cytomegalovirus, HCMV and MCMV, respectively. Abbreviations: i.p. = intraperitoneal injection, PFU = plaque-forming unit, MOI = multiplicity of infection, p\_adj = p value adjusted for multiple testing. #HCMV expression values are from the symptomatic samples compared to controls. ADAR2 values are from two different probes, ILMN\_1679797 and ILMN\_2319326, respectively.

### HCMV Gene Expression Analysis

In order to understand the effects of congenital HCMV infection, we used data available in BioProject PRJNA422858. The original analysis (Ouellette et al., 2020) used blood samples of infants with symptomatic and asymptomatic congenital HCMV infections and performed microarray analysis to obtain gene expression data. While this did not allow us to evaluate editing rates due to the lack of sequencing data and resultant inability to perform variant calling, it provided a robust sample to evaluate the changes in expression of ADAR genes caused by congenital HCMV infection.

Reanalyzing the data using the NCBI GEO2R tool (Barrett et al., 2013), both ADAR1 and ADAR2 were found to be significantly overexpressed in HCMV samples compared to controls. Specifically, both genes were overexpressed in symptomatic vs. control HCMV samples (p-adj of 7.31E-07, and 7.59E-05 and 4.00E-03 for ADAR1 p150 probe ILMN_1776777, ADAR2 probes ILMN_1679797 and ILMN_2319326, respectively), and asymptomatic vs. control HCMV samples (p-adj of 3.54E-04, and 6.34E-05 and 1.62E-02, for probes ILMN_1776777, ILMN_1679797 and ILMN_2319326, respectively) (Supplementary File 1). However, no differences were detected between asymptomatic and symptomatic HCMV samples (p-adj of 0.338, and 0.874 and 0.981 for probes ILMN_1776777, ILMN_1679797 and ILMN_2319326, respectively). In addition to this, Reactome pathway analysis (Jassal et al., 2020) was used to explore pathways experiencing differential gene expression. Both asymptomatic and symptomatic HCMV samples showed an enrichment in IFN signaling pathways among differentially expressed genes, including Interferon alpha/beta signaling linked to ADAR1 p150 expression (p-adj = 4.48E-04 and 0.001747396, and 0.005521773 and 0.010567466 for Interferon alpha/beta signaling (R-HSA-909733) and Interferon Signaling (RHSA-913531) pathways in asymptomatic and symptomatic samples, respectively). Full lists of differentially expressed genes and significantly overrepresented Reactome pathways are listed in the Supplementary File 1.

### MCMV RNA-seq Data Analysis

Next, we examined available RNA-seq data to investigate the effects of mouse cytomegalovirus (MCMV). The original dataset (Kveštak et al., 2021) performed RNA-seq of the microglia of 3 newborn mice infected with MCMV and 3 controls after 8 days of infection. We found that ADAR1 was overexpressed in infected samples (p = 0.007949), as shown in Figure 1A, while ADAR2/3 were not significantly changed (Figure 1B-C, respectively). In addition, 149 editing sites were detected at 69 unique genomic locations (Supplementary File 2A). These sites were disproportionately found in MCMV-infected samples, as shown in Figure 2A (p = 0.01129). Furthermore, these sites were not only more prevalent in MCMV samples, but were also edited at higher rates, as shown in Figure 2B (p = 0.02584). In addition, Supplementary File 3 shows that the range of editing rates is much wider in MCMV samples than controls. Site-specific analysis showed 21 sites with significantly increased editing in MCMV samples, with no sites seeing significant decreases in editing. This included three editing sites in the Lamp2 gene, a membrane glycoprotein involved in lysosomal functions, a site in the Kcnk6 potassium channel, and a site in Tmem9b, which enhances proinflammatory cytokine production (full list is available in Supplementary File 2B). Reactome pathway analysis of sites edited in MCMV infection showed overrepresentation (FDR < 0.1) among immune-related pathways (Supplementary File 4A). Notably, some of the editing targets, such as Cybb and Arhgdia, belong to multiple pathways, including neurologically relevant ones, such as RAC1 GTPase cycle (R-HSA-9013149), involved in neuronal development (de Curtis 2014).

**Figure 1.**
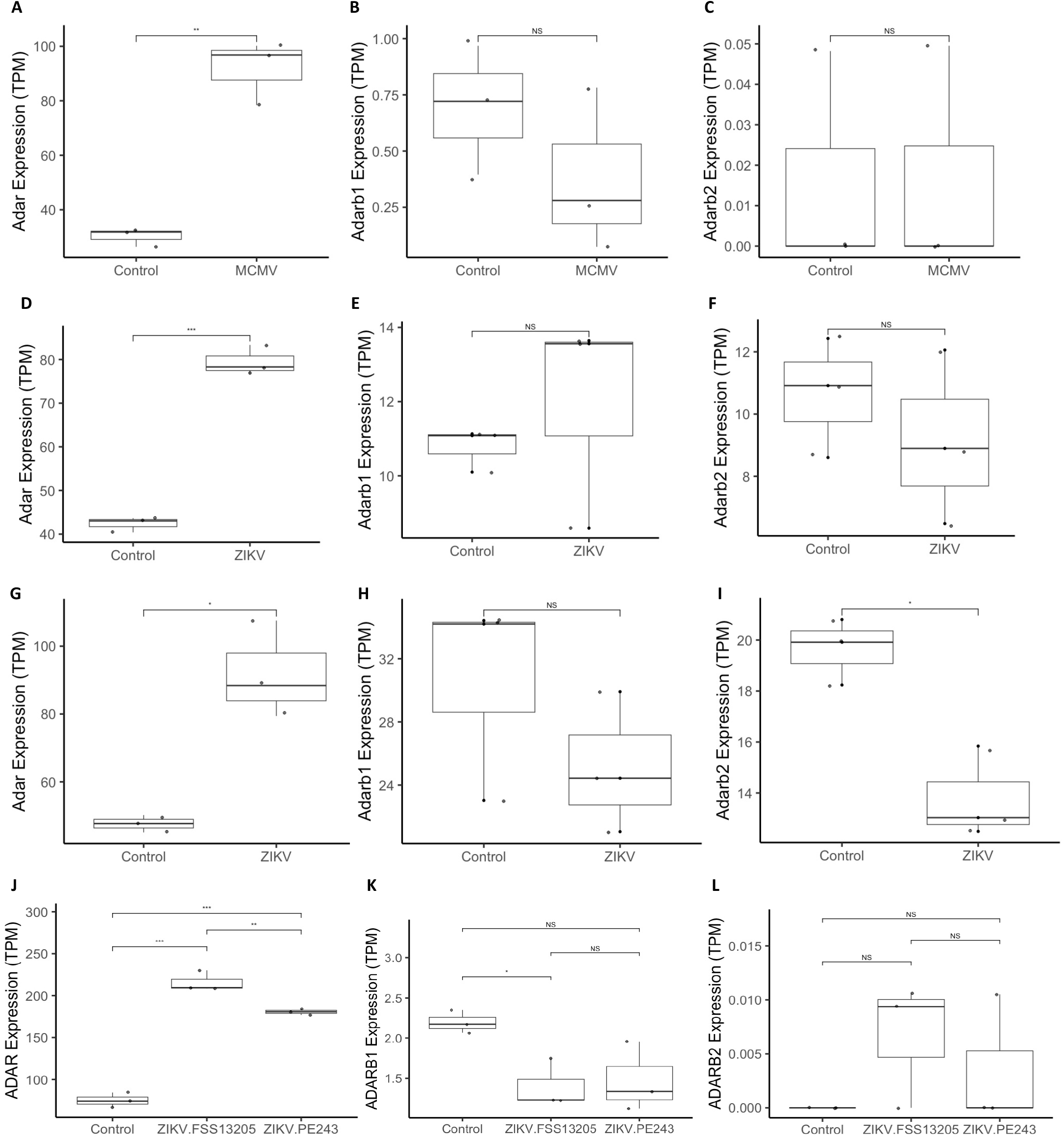
Figure 1. Box plots of ADAR1 (ADAR), ADAR2 (ADARb1) and ADAR3 (ADARb2) expression, in transcripts per million (TPM). Panels A, B and C depict expression differences between control vs. MCMV-infected mice samples. Only ADAR1 expression differed significantly between control and infected samples (p = 0.007949), but not that of ADAR2 or ADAR3 (p > 0.05). Panels D, E and F show expression in control vs. ZIKV-infected samples (PRJNA487357). Similar to MCMV infections, only ADAR1 expression differed significantly between control and infected samples (p = 0.0005574), but not that of ADAR2 or ADAR3 (p > 0.05). Panels G, H and I show expression in control vs. ZIKV-infected samples (PRJNA358758). Both ADAR1 and ADAR expression differed significantly between control and infected samples (p = 0.03035 and p=0.01266), albeit in different directions (over- and under-expressed, respectively), but not that of ADAR2 (p = 0.3086). Panels J, K and L show expression in control vs. ZIKV-infected human induced pluripotent neuroprogenitor stem cells (hiNPCs) samples (PRJNA551246). Infections with Cambodian (FSS13025) and Brazilian ZIKV (PE243) strains are shown separately. ADAR1 expression differed significantly between control and infected samples for both strains (padj = 1.81*10-5 and padj = 3.00*10-6, respectively), and was higher in FSS13025-infected samples compared to PE243V-infected ones (padj = 7.22*10-3). ADAR2 expression was significantly decreased in FSS13025-infected samples compared to controls (padj = 0.04909), but not in PE243V-infected ones (p = 0.06974). ADAR3 expression did not differ significantly between conditions.

**Figure 2.**
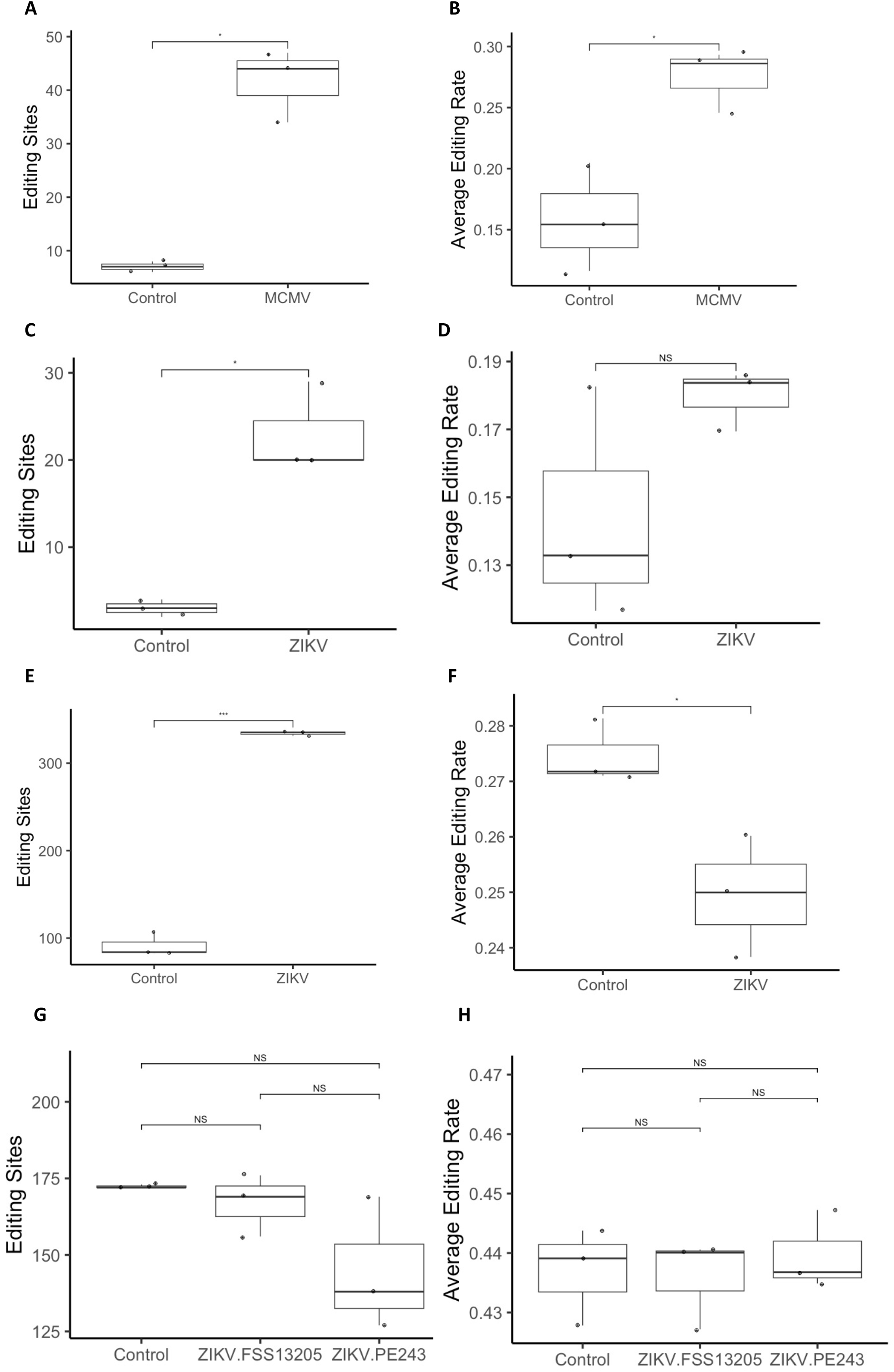
ADAR editing differences between control and infected samples, shown as the box plots of the number of editing sites and the average editing rate. Panels A and B show the editing sites and the editing rate, respectively, between control vs. MCMV-infected mice samples. Both differed significantly between control vs. MCMV-infected mice (p = 0.01129 and p = 0.02584, respectively). Panels C and D show the editing sites and the editing rate, respectively, between control vs. ZIKV-infected samples (PRJNA487357). The number of editing sites but not the average editing rates differed significantly between control vs. ZIKV-infected samples (p = 0.01877 versus p = 0.23096, respectively). Panels E and F show the editing sites and the editing rate, respectively, between control vs. ZIKV-infected samples (PRJNA358758). Both the number of editing sites and the average editing rates differed significantly between control vs. ZIKV-infected samples (p = 0.0007123 versus p = 0.03779, respectively). Unlike PRJNA487357-based dataset, here the editing rates appeared to be decreased in ZIKV-infected samples. Panels G and H show the editing sites and the editing rate, respectively, between ZIKV-infected human induced pluripotent neuroprogenitor stem cells (hiNPCs) samples (PRJNA551246). Infections with Cambodian (FSS13025) and Brazilian ZIKV (PE243) strains are shown separately. Unlike other ZIKV-infection examples, no significant differences were detected in the number of editing sites or average editing rate.

### Mouse ZIKV RNA-seq Data Analysis

We next wanted to investigate the effect of ZIKV on ADAR editing. To do this, we first used two RNA-seq datasets of ZIKV-infected mice. The first of these datasets (BioProject PRJNA487357), performed RNA-seq on total RNA from whole brains of 3 ZIKV infected and 3 control mouse embryos at E14.5 after infection at E10.5. ADAR1 was overexpressed in infected samples (p = 0.0005574) (Figure 1D). 78 editing sites were detected at 34 unique coordinates (Supplementary File 5A), disproportionately found in ZIKV samples, as shown in Figure 2C (p = 0.01877). However, unlike in MCMV, there was no statistically significant difference between editing rates of these sites (p = 0.23096) (Figure 2D). Despite this, Supplementary File 6 shows that the range of editing rates is somewhat increased in all ZIKV samples, even though the average editing rates are not significantly different.

A site-specific analysis showed that most sites experienced increased editing in ZIKV samples, and none with decreased editing (Supplementary File 5B). Interestingly, Gria2 was edited in both conditions with no significant difference in editing rate, while Blcap and Snhg11 showed higher levels of editing in ZIKV samples. Reactome pathway analysis of editing targets showed overrepresentation (FDR < 0.1) of several neurologically relevant pathways, including “unblocking of NMDA receptors, glutamate binding and activation,” “long-term potentiation,” “MECP2 regulates neuronal receptors and channels,” and “transcriptional regulation by MECP2”. The latter two pathways are noteworthy due to prominent role of MECP2 in modulating synaptic plasticity (Ebert and Greenberg 2013; Carstens et al., 2021) (Supplementary File 4B).

The second mouse ZIKV dataset (Chang et al., 2020) consisted of RNA-seq of the whole brains of 3 ZIKV SZ01 infected and 3 control embryonic mouse P3 brains infected at E15.5. ADAR was found to be overexpressed in ZIKV samples (p = 0.03035), while ADAR3 was underexpressed in ZIKV samples (0.01266) and ADAR2 was not significantly different (p = 0.3086) (Figure 1G-I, respectively). In addition, 1276 editing events were detected at 501 unique coordinates (Supplementary File 7A), disproportionately found in ZIKV samples (p = 0.0007123) (Figure 2F). Interestingly, in this dataset, average editing rates were decreased compared to controls (p = 0.03779) (Figure 2F, Supplementary File 8).

Numerous edited sites harbored significant differences in editing, including 11 sites with significantly decreased editing in ZIKV samples and 137 sites with significantly increased editing in ZIKV samples (Supplementary File 7B). This included 9 sites in exonic regions, including key neural genes such as Gria3, Grik5, and Nova1. While members of many of the same neural-related pathways were identified among edited targets, there were no overrepresented pathways that passed the FDR < 0.1 cut-off threshold in Reactome pathway analysis (Supplementary File 4C). Significantly, there was a high degree of overlap between the two mouse ZIKV datasets, as 9 of 12 coordinates and 9 of 10 genes with differential editing in PRJNA487357 were also differentially edited in PRJNA358758.

Due to the finding of differential editing in Nova1 and other splicing factors, we evaluated local splicing variations (LSVs) between infected and uninfected samples using MAJIQ and VOILA (Vaquero-Garcia et al., 2016). 936 LSVs with a change in percent spliced in (|dPSI|) > 0.2 and p < 0.05 in 650 genes (Figure 3; Supplementary File 9). Of these, 87 LSVs in 49 genes occurred in genes identified as Nova targets as per Zhang et al. (2010), indicating dysregulation of Nova RNA splicing. In addition, 25 LSVs were detected in 13 disease associated Nova targets, indicating the potential significance of this dysregulation. However, there is no way to determine whether this dysregulation is related to changes in Nova1 editing by ADAR.

**Figure 3.**
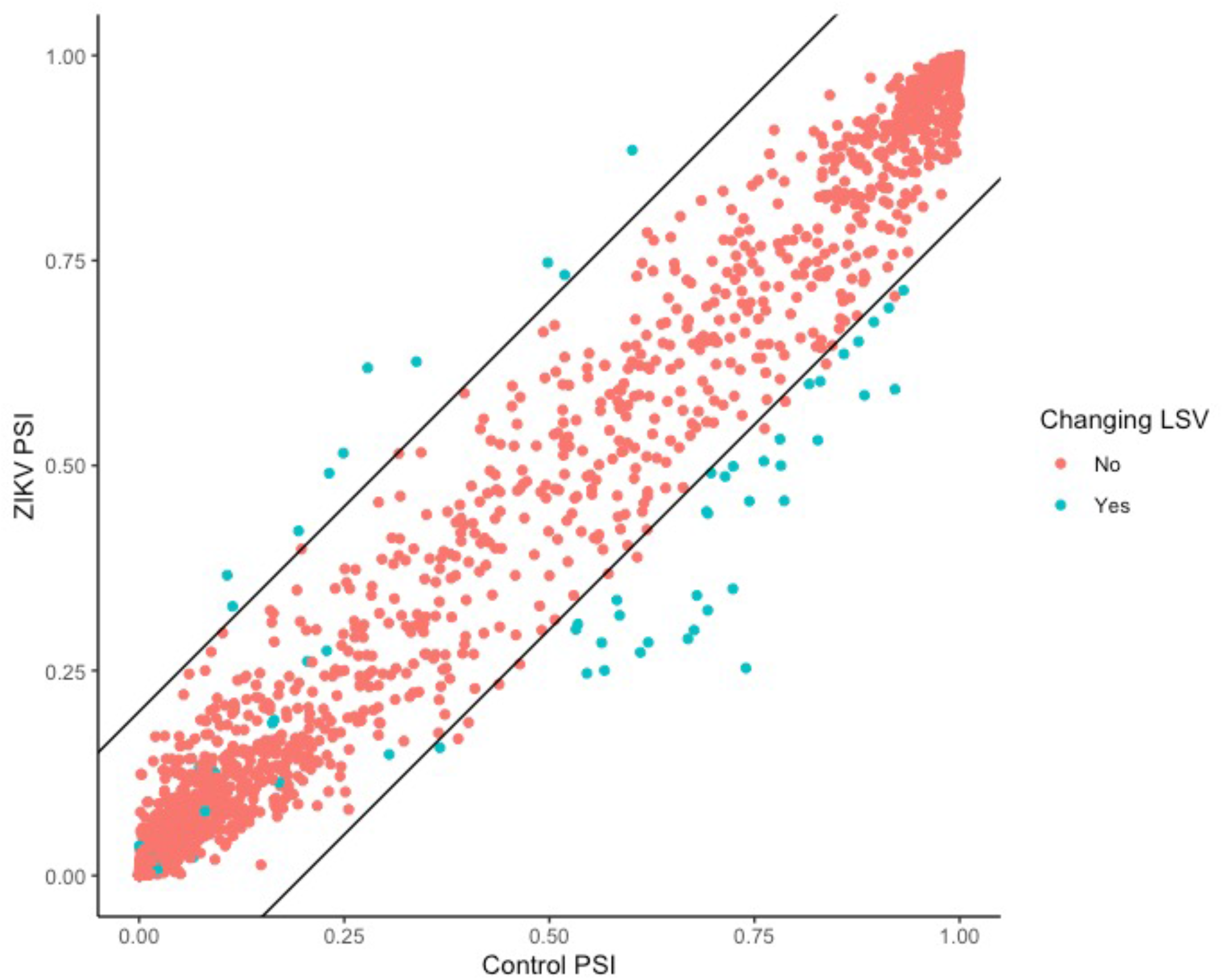
Scatterplot showing splicing changes between ZIKV and Control samples for PRJNA358758 for all LSVs in genes targeted by Nova1, as per Zhang et al., 2010. Significantly changing local splicing variations, LSVs ((|dPSI|) > 0.2 and p < 0.05), are shown in blue, with lines (slope = 1 and y intercept = -0.2/0.2) to show 0.2 change in percent spliced in (PSI) values.

### hiNPC ZIKV RNA-seq Data Analysis

We further analyzed a dataset of human induced pluripotent neuroprogenitor stem cells (hiNPCs), allowing us to look specifically at editing in NPCs and to get a perspective from human samples. The original study (Lima et al., 2019) took RNA-seq data from hiNPCs; 3 controls, 3 infected with the Cambodian strain of ZIKV (FSS13025), and 3 infected with the Brazilian strain of ZIKV (PE243). In these samples, 1355 editing sites were detected at 312 unique coordinates (Supplementary File 10A). ADAR1 expression was found to be higher in samples infected with both PE243V and FSS13025 ZIKV (padj = 1.81*10^-5^ and padj = 3.00*10^-6^), and was also found to be higher in samples infected with the FSS13025 strain compared to the PE243V strain (padj = 7.22*10^-3^) (Figure 1J). In addition, ADAR2 expression was significantly decreased in FSS13025 ZIKV samples compared to controls (padj = 0.04909), but not in PE243V ZIKV samples (p = 0.06974) (Figure 1K). No statistically significant differences were detected in ADAR3 expression between conditions; however, samples of both ZIKV strains had samples with elevated ADAR3 expression, whereas ADAR3 was not expressed in any control samples (Figure 1L). Unlike previous samples, no significant differences were detected in the number of editing sites or average editing rate; however, some ZIKV samples had a lower number of editing sites than the control samples (Figure 2G-H; Supplemental File 11).

Nonetheless, analysis of differences at specific editing sites found 9 sites with significantly different editing rates between conditions (Supplementary File 10B). Of these sites, 3 were increased and 2 were decreased in ZIKV FSS1302 compared to controls, one was increased in ZIKV PE243V compared to controls, one was increased in both ZIKV FSS1302 and PE243V compared to controls, and one was increased in ZIKV PE243V compared to ZIKV FSS1302. Specifically, all three editing sites which saw increased editing in ZIKV FSS1302 vs controls were in the DDX58 gene, which codes for the RIG-1 receptor. Additionally, B9D1, a gene involved with ciliogenesis, is almost completely edited in control samples, but only edited in one ZIKV PE243V sample and no ZIKV FSS1302 samples. This gene is implicated in the development of Meckel Syndrome, characterized by CNS developmental abnormalities, such as encephalocele. In both ZIKV FSS1302 and PE243V samples, an exonic site in the IFITM2 gene which is not edited in controls is highly edited (between 88% and 96%). IFITM2 is an IFN induced transmembrane gene responsible for restricting viral entry. The only site with significantly different editing between the two ZIKV strains occurred in the NCK2 gene. NCK2 is an adapter for receptor tyrosine kinases believed to be involved in cytoskeletal reorganization.

### Editing at miRNA Binding Sites

While a number of protein-recoding editing changes were observed, the majority of changing editing sites occurred in 3’ UTR regions. This is significant because of the 3’ UTR’s role in regulating gene translation and expression, including through the binding of miRNAs (Kuersten & Goodwin, 2003). Given previous reports that ADAR editing can affect miRNA binding (Peng et al., 2012; Wang I.X. et al., 2013; Wang Q. et al., 2013), we evaluated whether editing sites disrupted by congenital infection may have effects on gene expression by altering miRNA binding. After identifying differentially genes known to be targeted by miRNAs using TarBase (<u>Karagkouni</u> et al., 2018), SubmiRine (Maxwell et al., 2015) was used to evaluate differences in miRNA binding between unedited and edited transcripts. While no editing was found to alter miRNA binding in the MCMV or hiNPC ZIKV datasets, editing sites with potential links to changes in miRNA targeting were detected in 2 genes for the mouse ZIKV PRJNA487357 dataset and 26 genes for the mouse ZIKV PRJNA358758 dataset. Importantly, a number of these genes had links to neurological disease, development, and function (Supplemental File 12).

## Discussion

Congenital infections commonly cause neurodevelopmental abnormalities through poorly understood mechanisms, often implicating the immune response. Here, we raised the question of whether the IFN-driven induction of ADAR RNA editing enzyme expression affects editing of host transcriptome, and whether such editing dysregulation may be related to the neurodevelopmental abnormalities caused by congenital infection (Piontkivska et al. 2019). Our RNA-seq data analyses provide evidence for changes in ADAR expression and editing caused by congenital infection by CMV and ZIKV in mice, and to a lesser degree, ZIKV infection of hiNPCs. Crucially, our findings also show changes in editing in genes relevant to neural development and function, providing a potential link to the neurodevelopmental abnormalities caused by congenital infection. In mouse datasets, most of these sites saw increased editing, but a small number of sites also saw decreased editing rates in infected samples. The dataset of ZIKV infected hiNPCs also saw a greater number of sites with increased editing, but differences in dysregulated editing sites were observed between ZIKV strains. There was significant overlap between sites differentially edited in different mouse datasets, particularly between the two ZIKV datasets, which further provides support for the reliability of these findings.

One of the most significant editing sites that showed differential editing was the AMPA glutamate receptor subunit GRIA3 R/G recoding site, which was consistently underedited in ZIKV-infected samples for the PRJNA358758 dataset. Normally, editing of this site displays a gradual increase in editing through development (Wahlstedt et al., 2009). The resulting R/G recoding causes faster recovery from desensitization, allowing quicker responses to impulses (Lomeli et al. 1994). This finding reinforces previous evidence of altered GRIA3 editing caused by ZIKV infection (Piontkivska et al., 2019). Another significant neurological target that showed differential editing (increased) in the PRJNA358758 dataset was the GRIK5 K/R recoding site. GRIK5 is another excitatory glutamate receptor for which reduced expression has been associated with eye and vascular disease (Unlu et al., 2019), and whose variants have been associated with neurological disorders (Koromina et al., 2019). While this gene has been shown to be consistently edited, the effect of K/R recoding on protein function neurological phenotype has not been fully elucidated (Danecek et al., 2012; Huntley et al., 2016).

Another one of the primary neurological genes found here was Calm1, edited in PRJNA487357 and PRJNA487357 mouse ZIKV datasets. Calm1 encodes a protein of the calmodulin family: these proteins are Ca^2+^ sensors, acting as secondary messengers with a wide range of functions. For example, calmodulin plays a role in regulating long-term potentiation (LTP), a mechanism for synaptic plasticity relevant to learning and memory (Lledo et al., 1995). Calmodulin can also affect smooth muscle contraction (Martinsen et al., 2014), which has implications for numerous important physiological processes. Editing of Calm1 is not the only part of the Ca^2+^ signaling pathway impacted by differential editing in our data. A downstream effector of Calm1, calmodulin-dependent kinase IV (Camk4), also saw altered editing in the PRJNA358758 mouse ZIKV dataset. This gene is expressed in lymphocytes and neurons (Anderson et al., 1998) and has a diverse array of immune and neurological functions (Naz et al., 2020). In the brain, Camk4 regulates synaptic excitation (Joseph & Turrigiano, 2017), memory formation (Fukushima et al., 2008), protects neurons from apoptosis (See et al., 2001), and has been implicated in neurodevelopmental disease (Zech et al., 2018). It also regulates differentiation into Th17 cells and IL-17 production, and plays a role in regulating dendritic cell lifespan (Koga & Kawakami, 2017). Another calmodulin binding protein which saw dysregulated editing is Map6 (a.k.a. STOP), which was differentially edited in the PRJNA358758 mouse ZIKV dataset. Map6 stabilizes microtubules and is targeted to the axons of polarizing neurons, playing an important role in axon maturation and the establishment of polarity (Tortosa et al., 2017). In addition to this, Map6 plays an important role in Sema3E signaling and is required for proper axonal extension of subicular neurons and proper development of brain regions such as the fornix (Deloulme et al., 2015). MAP6-KO mice have defects in neurotransmission and synapse formation, and display a range of cognitive and behavioral impairments, including pathology similar to schizophrenia (Andrieux et al., 2002; Benardais et al., 2010; Brun et al., 2005; Deloulme et al., 2015; Fournet et al., 2010; Gimenez et al., 2017; Volle et al., 2013).

Interestingly, differential editing occurred in several genes responsible for regulation of mRNA splicing. The most significant of these was the Nova1 S/G recoding site (overedited in PRJNA358758), but also included 3’ UTR editing sites in genes such as spliceosome subcomponent Sf3b2 (overedited in all mouse datasets) and RNA binding protein Celf1 (overedited in PRJNA358758). Dynamic regulation of alternative splicing plays an important role in brain development, and dysregulation of this process has implications for neurodevelopmental disorders (Han et al., 2022; Porter et al., 2018; Raj & Blencowe, 2015). Specifically, Nova1 is a neurologically expressed RNA binding protein responsible for regulating alternative splicing. In addition, alternative splicing of cryptic exons by Nova regulates RNA abundance of a number of genes with synaptic functions through nonsense mediated decay (Porter et al., 2018; Raj & Blencowe, 2015). Nova1-knockout mice experience neuronal apoptosis in the brain stem and spinal cord, followed by motor dysfunction and postnatal death (Porter et al., 2018; Raj & Blencowe, 2015). ADAR editing and S/G recoding of Nova1 is highly evolutionarily conserved and its effects are relatively well documented (Irimia et al., 2012). This editing site is dynamically regulated with increasing levels through brain development, and has been shown to increase Nova1 protein stability by protecting against proteasomal degradation (Irimia et al., 2012) In doing so, dysregulation of this editing site has been implicated in neurological disease in both humans and mice (Karagianni et al., 2022). Indeed, in PRJNA358758 where Nova1 S/G editing was found to be dysregulated, alternative splicing of a Nova targets was detected, including in 13 disease-associated genes.

Another area of interest is the potential disruption of epigenetic genome regulation. Phc2 (also known as Mhp2), a component of the class II Polycomb gene (PcG) complex as well as the Polycomb repressive complex 1 (PRC1) (Isono et al., 2013) has differential editing in the 3’ UTR region, which may alter miRNA targeting, in the PRJNA358758 and PRJNA487357 mouse ZIKV datasets. Disruption of class II PcG genes alters the expression of Hox cluster genes in the paraxial mesoderm and neural tube and causes axial skeleton malformations (Isono et al., 2005). Phc2 is expressed in NPCs (Kim et al., 2005) and represses the expression of neurogenic genes during later stages (Tsuboi et al., 2018). This illustrates an interesting possibility by which editing dysregulation could disrupt neurodevelopment.

The analysis of hiNPC data revealed a different set of differentially edited sites. Two differentially edited genes are associated with neurodevelopmental disorders. An intronic site in the NCK2 was overedited in cells infected with ZIKV PE243V compared to the FSS1302 strain, while another intronic site in the B9D1 gene was unedited in the ZIKV FSS1302 strain, but highly edited in controls. NCK2 is a tyrosine kinase adaptor responsible for regulating cytoskeleton organization, and may thusly contribute to the formation of proper neuronal connections (Fawcett et al., 2007; Pasquale, 2008; Wegmeyer et al., 2007). B9D1 is important for ciliogenesis and has been implicated in ciliopathies including Meckel syndrome (Dowdle et al., 2011; Hopp et al., 2011), characterized by renal cystic dysplasia and CNS defects, and Joubert syndrome (Bachmann-Gagescu et al., 2015; Kroes et al., 2015; Romani et al., 2014), characterized by cerebellar and brainstem malformation.

However, a number of potentially significant editing changes were also observed in immune genes for the hiNPC dataset. First, an exonic V/A recoding site in the IFITM2 gene was overedited in cells infected with both strains of ZIKV compared to controls. IFITM2 is an interferon-stimulated transmembrane protein which restricts viral membrane fusion and entry of many viruses including ZIKV (Li et al., 2013; Savidis et al., 2016; Brass et al., 2010). While the impact of this recoding site on IFITM2 protein function is currently unknown, this might mean that changes in ADAR editing could influence the efficiency of ZIKV entry. Additionally, three sites in the 3’ UTR of DDX58 (RIG1) were overedited in the ZIKV FSS1302 strain compared to controls. RIG1 is an RNA helicase and dsRNA sensor critical for inducing the antiviral type 1 IFN response. Taken together, these could indicate that ADAR editing could modulate other aspects of the interferon response.

Aside from these, many other sites were impacted by differential ADAR editing, with some notable highlights described in Table 2. Future studies could use sites listed here as candidates to validate the neurodevelopmental effects of virus-induced editing dysregulation. Many neurological genes with dysregulated editing could contribute to death of neuronal cells, disrupt the formation of neuronal connections or lead to altered synaptic transmission. Disruption of other genes such as those regulating splicing programs or epigenetic gene silencing could alter regulatory programs in NPCs at a crucial point in their development. In addition, the immunological targets with differential editing could lead to a dysregulated immune response, which could have adverse effects through increased susceptibility to viral infection or immune mediated damage or alterations to neurons or NPCs.

**Table 2.** List of genes with significant editing dysregulation, namely, observed differential editing in exonic regions with effects on protein-coding regions, in neurologically or immune relevant pathways, or in multiple samples. Parenthetical numbers next to a dataset for a given gene specify that >1 site was differentially edited in that gene/dataset.

| Gene | Dataset(s) | Genic location of edited site/Effect | Function/significance of edited gene |
| --- | --- | --- | --- |
| **Neurological:** |  |  |  |
| Gria3 | PRJNA358758 | Exon (nonsyn) | AMPA glutamate receptor subunit, editing allows faster recovery from desensitization (Lomeli et al. 1994) |
| Grik5 | PRJNA358758 | Exon (nonsyn) | Kainate glutamate receptor subunit associated with psychiatric, eye, and vascular diseases (Unlu et al., 2019; Koromina et al., 2019) |
| Calm1 | PRJNA358758, PRJNA487357 | 3' UTR | Ca<sup>2+</sup>-binding messenger impacting numerous neurological functions (Solà et al., 2001; DeLorenzo, 1982; Lledo et al., 1995) |
| Camk4 | PRJNA358758 | 3' UTR | Signal transducer downstream of Calmodulin with important and diverse neurological and immune functions (Naz et al., 2020) |
| Map6 | PRJNA358758 | 3' UTR | Calmodulin binding protein regulating microtubule stability, important for proper axon development, disruption leads to issues with neurotransmission and synapse formation, as well as cognitive/behavioral deficits (Tortosa et al., 2017; Deloulme et al., 2015; Brun et al., 2005; Benardais et al., 2010; Fournet et al., 2010; Volle et al., 2013; Andrieux et al., 2002; Gimenez et al., 2017) |
| Selenot | PRJNA358758 (2),<br>PRJNA487357 | 3' UTR | Thioredoxin-like oxidoreductase, neuroprotective, highly expressed during brain development, KO affects brain structure through neuron loss and causes behavioral changes, important role in brain development (Anouar et al., 2018) |
| Sgpl1 | PRJNA358758,<br>PRJEB38849 | 3' UTR | sphingosine phosphate lyase, regulates neuronal autophagy (Mitroi et al., 2016), microglial autophagy and inflammation (Karunakaran et al., 2019), mutations linked with neurological pathologies (Alam et al., 2020; Martin et al., 2020; Bamborschke et al., 2018) |
| Arhgdia | PRJNA358758,<br>PRJEB38849 (2) | 3' UTR | Regulator of Rho GTPase signaling, regulates cell proliferation and migration, underexpression promotes glioma progression (Lu et al., 2016; Lin et al., 2016); Rho GTPase signaling plays an important role in neurodevelopment and dysregulation may lead to neurological disorders (Govek et al., 2005) |
| NCK2 | PRJNA551246 | Intron | tyrosine kinase adaptor protein regulating cytoskeleton organization and formation of neuronal connections (Pasquale, 2008; Fawcett et al., 2007; Wegmeyer et al., 2007) |
| **Immune:** |  |  |  |
| Lgals3 | PRJNA358758 | Exon (nonsyn) | Galectin with affinity for beta-galactosides, can regulate adhesion/inflammation of immune cells, can affect cell growth/differentiation, including immune cell/neurite growth, and acts as a splicing factor along with other functions (Dumic et al., 2006; Pesheva et al., 1998; Dagher et al., 1995) |
| Tapbp | PRJNA358758 (3),<br>PRJNA487357 (2),<br>PRJEB38849 (2) | 3' UTR | Mediates interaction between MHC1 and TAP to allow loading of antigenic peptides (Zhang & Williams, 2006) |
| Xbp1 | PRJNA358758,<br>PRJEB38849 | 3' UTR | Transcription factor regulating immune and UPR functions, also with links to neurodegenerative disease (Acosta-Alvear et al., 2007) |
| Ube2d3 | PRJNA358758 | 3' UTR | E2 ubiquitin ligase, involved in RIG-1 activation (Shi et al., 2017) |
| RIG1 | PRJNA551246 (3) | 3' UTR | dsRNA sensor responsible for activating innate antiviral type 1 interferon response (Kell & Gale, 2015) |
| IFITM2 | PRJNA551246 | Exon (nonsyn) | IFN-stimulated antiviral restriction factor (Li et al., 2013; Savidis et al., 2016; Brass et al., 2010) |
| **Other:** |  |  |  |
| Ucp2 | PRJNA358758 | Exon (nonsyn) | Mitochondrial uncoupling protein (proton leak), attenuates mitochondrial ROS production (Brand & Esteves, 2005). Reduced expression is linked to altered differentiation of NPCs and has a significant role in brain development (Wang et al., 2022; Simon-Areces et al., 2012) |
| Ogdh | PRJNA358758 | Exon (stoploss) | 2-oxoglutarate dehydrogenase complex subunit, some evidence of links to neurological disease (Szabo et al., 1994; Yap et al., 2020) |
| Azin1 | PRJNA358758 | Exon (nonsyn) | Antizyme inhibitor regulating intracellular polyamine levels, ADAR editing of this gene is linked to development of a number of cancers, including colorectal, non-small-cell lung, gastric, and hepatocellular cancers (Shigeyasu et al., 2018; Takeda et al., 2019; Wei et al., 2022; Hu et al., 2017; Okugawa et al., 2018; Chen et al., 2013), as well as to hematopoietic stem cell differentiation (Wang et al., 2021) |
| Nova1 | PRJNA358758 | Exon (nonsyn) | RBP regulating splicing and degradation of a number of genes with important neurological functions through brain development: editing is dynamically regulated during development and increases protein stability (Raj & Blencowe, 2015; Porter et al., 2018; Irimia et al., 2012) |
| Celf1 | PRJNA358758 | 3' UTR | RBP regulating splicing during brain development (Ladd, 2013) |
| Sf3b2 | PRJNA358758,<br>PRJNA487357,<br>PRJEB38849 | 3' UTR | Splicing factor, U2 snRNP component, variants associated with craniofacial microsomia (Timberlake et al., 2021) |
| Sept2 | PRJNA358758,<br>PRJNA487357,<br>PRJEB38849 | 3' UTR | Cytoskeletal GTP-binding filament-forming protein (Kremer et al., 2007) Sept2 is expressed in the brain, and neurological functions include regulation of astrocyte glutamate uptake (Kinoshita et al., 2004) |
| Lamp2 | PRJNA358758 (3),<br>PRJEB38849 (3) | 3' UTR | Lysosome membrane glycoprotein, plays a role in autophagy regulation (Eskelinen, 2006) |
| Phc2 | PRJNA358758,<br>PRJNA487357 | 3' UTR | Component of class II PcG complex and PRC1, contributes to epigenetic regulation of gene expression, including neurogenic genes during brain development (Isono et al., 2013; Kim et al., 2004; Tsuboi et al., 2018) |
| H19 | PRJNA358758,<br>PRJNA487357 | lncRNA | lncRNA which functions as a tumor suppressor and regulates growth during embryonic development (Gabory et al., 2006; Gabory et al., 2010) |
| Gpx3 | PRJNA358758,<br>PRJEB38849 | 3' UTR | Glutathione peroxidase, reduces hydrogen peroxide to prevent oxidative damage (Takahashi et al., 1987) |
| Fam49b<br>(a.k.a. Cyrib) | PRJNA358758,<br>PRJEB38849 | Exon (syn) | Interacts with Rac GTPase, functions include mitochondrial ROS suppression, modulating cytoskeleton organization, and inhibition of T cell activation (Shang et al., 2018; Chattaragada et al., 2017) |
| Tmem50b | PRJNA358758,<br>PRJEB38849 | 3' UTR | Transmembrane protein, ER localization, may contribute to proper brain development, with dysregulation leading to Down syndrome-related phenotypes (Moldrich et al. 2008) |
| Cap1 | PRJNA358758 | 3' UTR | Regulates cytoskeleton organization, adhesion, cAMP signaling (Zhang et al., 2013). Also plays a role in regulating neuron differentiation (Schneider et al., 2021a; Schneider et al., 2021b) |

As shown by the changes in ADAR editing and expression summarized in Tables 1 and 2, there is significant heterogeneity in the changes observed in the different datasets we analyzed here. Only the MCMV dataset saw increased average editing rates, and one mouse ZIKV dataset saw decreased average editing rates, even though the total number of editing sites was increased in every dataset except the hiNPC ZIKV dataset. In addition, while all datasets saw increased ADAR1 expression during infection, one mouse ZIKV dataset also saw decreases in ADAR3 expression, and the hiNPC ZIKV dataset saw decreases in ADAR2 expression. This decrease in ADAR2 expression in ZIKV-infected samples could help explain the lack of global editing changes that were seen. It may be the case that in these samples, increased editing from overexpressed ADAR1 was compensated by decreased editing from underexpressed ADAR2. This may also be related to autoregulation of ADAR2 expression, as rodent ADAR2 has the ability to edit its own pre-mRNA, creating an alternative splice site that leads to a frameshift and subsequent decrease in ADAR2 expression (Rueter et al., 1999).

In addition, it is possible that differences in experimental procedures used in these studies may be responsible for observed differences between datasets. For example, differences in viral delivery method and viral load (as summarized in Table 1) could lead to differences in immune responses or patterns of infection that affect ADAR editing. In addition, differences in the developmental stages at which infection and sequencing occurred could further contribute to differences between samples. Given the dynamic regulation of ADAR editing through development, as well as general changes in transcriptome composition over time, interventions at different stages may have different effects on editing. The length of time between infection and sample collection could also impact editing changes observed, especially in light of the work of Tsivion-Visbord et al. (2020) demonstrating the transient nature of MIA-induced editing changes. Another critical factor that may result in editing variations is biological sex, which can result in significant differences in editing patterns and was not examined here.

Aside from these factors, the observed differences in editing effects are consistent with previous studies showing that a variety of host- and virus-specific factors can influence changes in ADAR expression and editing induced by viral infection (Cao et al., 2018; Piontkivska et al., 2021). In previous studies, viral infection has been shown to have diverse effects on host editing, including editing increases during measles virus (Suspène et al., 2008) infection and no editing changes for Reovirus (Hood et al., 2014). Piontkivska et al. (2019) subsequently found that changes in editing induced by ZIKV in human stem cell cultures varied based on cell line, which may be in part related to differences in innate immune pathways and activation of ADAR1 p150 expression. In addition, Cao et al. (2018) found remarkable variance in effects on host ADAR editing even between different strains of influenza A virus (IAV) with additional variance between viral hosts. Our analysis parallels the previous literature, with differences in editing based on both virus (MCMV vs. ZIKV) and host (ZIKV in mice vs. hiNPCs), and with differences in editing and ADAR expression between strains of ZIKV. Notably, both mouse ZIKV datasets displayed similar patterns in editing, with 9 out of 12 coordinates and 9 out of 10 genes with differential editing in the PRJNA487357 dataset also seeing differential editing in the PRJNA358758 dataset, a level of overlap much greater than between any other datasets. This suggests a role for an unknown set of highly sensitive factors in both host and virus which regulate changes in host transcriptome editing in response to viral infection.

After considering changes in editing, it is important to biologically contextualize the results. First, the MCMV data provides evidence for editing dysregulation in microglia during congenital infection, though we have no data on neurons or NPCs for this virus. In addition to this, we found that ADAR expression is upregulated in congenital HCMV infections. The relationship between ADAR editing and expression seen in the MCMV samples may indicate that overexpression of ADAR1 and ADAR2 seen in HCMV implies increased editing as well. However, this may not be the case for all genes and/or edited sites, given the complex relationship between ADAR expression and editing, including host/virus-specific factors (Piontkivska et al., 2021). Further evidence would be needed to determine the specifics of editing dysregulation during HCMV infection, given that editing patterns vary by cell type, specific transcript, and sex of organism, often involving yet-to-be-determined regulatory mechanisms.

Next, the ZIKV mouse data gives evidence of significant editing changes that may disrupt neural function during development. However, for both of these datasets, bulk brain tissue sequencing was used, which obscures insights into the potential brain region and cell type specific regulation of editing (Cuddleston et al., 2022; Lundin et al., 2020). The potential pitfalls of inferring editing changes from whole brain sequencing are shown by Gal-Mark et al. (2017), that found that supposed changes in editing following spinal cord injury inferred by others such as Barbon et al. (2010) were simply due to decreased neuron density, as neurons have different editing rates than other CNS cell populations. The limited evidence of editing dysregulation in the hiNPC dataset gives greater insight into editing changes in humans, as well as specifically in the NPC population, though it is limited by being an *in vitro* cell culture model. Ideally, future research would include sequencing of specific cell type populations to delineate editing changes in NPCs and effects on neural development, as well as immune cells to determine effects of ADAR editing on the immune response are also of potential significance. Additionally, given that editing patterns have been shown to vary based on biological sex (Silvestris et al., 2019), it is significant that none of the datasets here included this as a variable. Future studies should fill this gap by examining how editing is differentially affected in both males and females to determine if there are biologically significant differences, and whether this has implications for differential development of neurodevelopmental symptoms.

The differences between the Brazilian (FSS13025) and Cambodian (PE243V) ZIKV strains are also of potential significance, and may be related to differences in immune responses induced by different ZIKV strains (Hernandez-Sarmiento et al., 2023). A higher number of editing sites were dysregulated compared to controls in the Cambodian strain compared to the Brazilian strain, and one site in the NCK2 gene, responsible for cytoskeleton regulation and formation of neuronal connections (Fawcett et al., 2007; Pasquale, 2008; Wegmeyer et al., 2007). While the observation of congenital abnormalities with ZIKV infection were first observed during the Brazilian epidemic, it is unclear whether this is due to differences in the effects of the virus or due to differences in reporting. Majumder et al. (2018) analyzed data on congenital birth defects from a West African hospital with endemic ZIKV, and found seasonal patterns in birth defects consistent with the spread of ZIKV by mosquitoes. Additional research has shown associations between infections with Asian ZIKV strains and congenital symptoms similar to those seen in Brazil (Chu et al., 2018). With the relative rarity of the most severe symptoms such as microcephaly, and the less obvious nature of other CZS symptoms, it could be the case that other ZIKV strains cause similar congenital abnormalities to the Brazilian strain which have simply gone unnoticed and unreported. Consistent with other studies (Esser-Nobis et al., 2019; Lima et al., 2019), we have found here that there are significant biological differences between different ZIKV strains. However, it is unclear whether or not these differences are associated with differences in symptoms of congenital infection. Future research should explore the exact biological implications of these differences, such as the differential editing sites found here.

There are a number of other technical limitations to our findings from the data analyzed here. First, it is worth noting that the number of editing sites found here was relatively small for some datasets, potentially owing to a lack of depth in sequencing. Sequencing depth ranged from 16-25 million reads in PRJEB38849 MCMV samples and 21-33 million reads for PRJNA487357 ZIKV samples, whereas for paired end Illumina sequencing, 80-100 million reads is generally desirable for RNA editing detection (Diroma et al., 2019). Future research using high depth sequencing would be helpful to fully illuminate the range of editing dysregulation caused by congenital infection. The benefits of this approach are illustrated by Tsivion-Visbord et al.’s (2020) analysis of MIA-induced RNA editing changes, where experiment repetition with 220 million read coverage revealed more robust changes in RNA editing and new dysregulated editing sites. Additionally, all RNA-seq datasets used here had three samples per condition, which limits the extent to which these findings can be extrapolated to the broader population.

Another serious limitation of our study is the inability to examine potential editing independent effects of changes in ADAR expression. ADAR can have many cellular effects through direct interactions with other proteins and competitive binding to RNA. First, in the immune response, ADAR1 can bind to and inhibit the activity of PKR, an IFN-induced gene which plays a role blocking translation as part of the antiviral immune response (Gélinas et al., 2011; Pfaller et al., 2021). In addition, ADAR affects a number of RNA processing pathways independent of editing. For example, ADAR1 can form a complex with miRNA processing protein Dicer to promote the rate of pre-miRNA cleavage and formation/loading of the RISC complex (Nishikura, 2016; Ota et al., 2013; Vesely et al., 2012; Yoshida et al., 2021). ADAR2 can also regulate splicing independent of editing by competing with U2AF65 for 3′ splice site binding. Other editing-independent effects of ADAR include regulation of gene expression through interactions with HuR (Wang I.X. et al., 2013) and NF90 (Nie et al., 2005). Given that ADAR expression was drastically altered in samples with viral infections, many of these processes may be altered in ways that cannot be linked to editing. Future research should explore this possibility.

The evidence presented here is consistent with the hypothesis that congenital CMV and ZIKV infection induces changes in ADAR editing, which in turn disrupts brain development. However, a causal link between virus-induced RNA editing dysregulation of specific transcripts and neurodevelopmental symptoms remains to be established. A useful first step here would be to test the effects of congenital infection with knock-downs or inhibition of ADAR and/or IFNs at different times during fetal development. This could help determine what role, if any, ADAR has in the development of specific neurodevelopmental symptoms. It may also be useful to test the effects of congenital infection in mice/cells with editing-inactive ADAR enzymes. This could help delineate what effects of ADAR are due to editing, and which may be due to ADAR interactions with other proteins. Following from this, the specific editing sites/interactions that result in neurodevelopmental abnormalities could be probed. This may involve testing the effects of specific editing sites by investigating brain development in the presence of different RNA variants or different combinations of RNA variants, or investigating the activity of ADAR binding partners. These factors should be tested in different cell types and brain regions, and at different developmental stages. This would give us a much more granular, mechanistic understanding of the relevance of ADAR editing during congenital infection in the brain.

## Conclusions

Overall, our results support the hypothesis that congenital viral infections by ZIKV and CMV induce expression of ADAR RNA editing enzymes and disrupt normal host transcriptome regulation during brain development. This has significant implications for understanding the mechanisms behind the pathogenesis of neurodevelopmental sequelae of congenital viral infections. We also lay out further experimentation that is necessary to confirm this hypothesis and to give a full mechanistic understanding of the effects of ADAR during congenital infection. We also suggest that future research should elucidate editing patterns caused by congenital infection of other viruses, given the highly virus-dependent nature of editing changes. This would be useful to inform potential novel treatment pathways, and to better understand the risks viral infection can pose to the developing brain.

## Methods

### BioProject Datasets

Data from BioProjects PRJNA422858, PRJEB38849, PRJNA487357, PRJNA358758, and PRJNA551246 were used. BioProject PRJNA422858 contains microarray expression data from blood samples of infants with asymptomatic and symptomatic human CMV (HCMV) infections, and healthy controls. BioProject PRJEB38849 contains RNA-seq data from newborn mouse microglia, 3 with mouse CMV (MCMV) infection, and 3 controls. BioProject PRJNA487357 and PRJNA358758 contain RNA-seq data from fetal mouse brain tissues, 3 infected with ZIKV, and 3 controls. And PRJNA551246 contains human induced pluripotent neuroprogenitor stem cells (hiNPCs), 3 infected with the Cambodian strain of ZIKV (FSS13025), and 3 infected with the Brazilian strain of ZIKV (PE243V), and 3 controls.

### Microarray data analysis

For BioProject dataset PRJNA422858/GEO dataset GSE108211 (HCMV), GEO2R (https://www.ncbi.nlm.nih.gov/geo/geo2r/) (Barrett et al., 2013) was used to evaluate differential gene expression, corrected for multiple hypothesis testing (FDR, false discovery rate) with the Benjamini–Hochberg procedure (Benjamini and Hochberg, 1995), quantile normalization and log2 transformation as implemented in GEO2R, with a specific focus on ADAR expression. Reactome (Jassal et al., 2020) was then used to identify overrepresentation of differentially expressed immune pathways that may be related to the changes in ADAR expression. However, because the dataset only contains microarray gene expression data, the extent of editing could not be evaluated for these samples.

### RNA-seq data analysis: variant calling and identification of editing sites

Next, the RNA-seq datasets were analyzed for ADAR expression and editing. For these samples, variant frequency counts produced by the Automated Isoform Diversity Detector (AIDD) pipeline (Plonski et al. 2020) were used to evaluate the extent of RNA editing. Briefly, AIDD used HISAT2 (Kim et al., 2015) for genome alignment with the GRCm38/mm10 reference genome for mouse samples and the hg37 reference genome for human samples, followed by StringTie (Pertea et al., 2015) for genome assembly and transcript counting as transcripts per million (TPM). GATK HaplotypeCaller (DePristo et al., 2011) was then used for variant calling to identify ADAR edited sites. Bam-readcount (Khanna et al. 2022) was used to determine the number of individual bases observed at each potentially edited site. Editing sites were defined as those variants that occur in the REDIportal database V2.0 (Mansi et al., 2021) with a reference of A or T (to identify A-to-G sites, or T-to-C as the nucleotide change would be interpreted on the opposite strand), with greater than 3 total reads, and with an editing rate (defined as the percent of G reads for an A reference sites or C reads for a T reference site) greater than 0.01, less than 0.99, and not between 0.49 and 0.51 (to remove potential noise, homozygous genomic variants, and heterozygous genomic variants, respectively).

### Statistical analysis of RNA-seq editing and expression data

Student’s t-test performed on TPM values were used to determine differential expression of ADAR enzymes, corrected for multiple hypothesis testing (FDR). Similarly, the average editing rate and number of editing sites in each sample were compared between infection and control conditions. In addition, for editing sites in both infected and control samples, editing rates were compared between conditions. T-tests (pairwise in the case of hiNPC data) were performed on editing rates, and corrected for multiple hypothesis testing with the Benjamini–Hochberg procedure. Significance was determined with a threshold of 0.05. Finally, Reactome pathway overrepresentation analysis was performed for editing sites with significant differences in editing.

### Analysis of the Effects of Editing on miRNA Binding

Sequences of 3’ UTRs with differentially edited sites were obtained from Ensembl Biomart (Cunningham et al., 2022), and editing coordinates were converted to their GRCm39 equivalents using CrossMap (Zhao et al., 2014). This step was used to generate edited versus wild type sequences for each edited gene/site with available sequences for all transcript isoforms containing the given coordinate. TarBase (Karagkouni et al., 2018) was used to identify genes known to be targeted by miRNAs, and miRNA sequences were obtained from miRBase (Kozomara & Griffiths-Jones et al., 2011). Finally, these sequences were used to find differences in miRNA targeting between edited and unedited transcripts using SubmiRine (Maxwell et al., 2015).

### Analysis of Differential Splicing

MAJIQ and VOILA software packages (Vaquero-Garcia et al., 2016) were used to assess changes in alternative splicing between infected and control samples in PRJNA358758 (HCMV). MAJIQ builder constructed splice graphs, and MAJIQ quantifier was used to quantify percent spliced in (PSI) and delta PSI (dPSI) of local splicing variations (LSVs). VOILA was used to output LSVs with |dPSI| > 0.2 with p < 0.05.

## List of abbreviations

ADAR: Adenosine Deaminases Acting on RNA
CMV: cytomegalovirus
CNS: central nervous system
CZS: congenital Zika syndrome
FDR: false discovery rate
HCVM: human cytomegalovirus
hiNPCs: human induced pluripotent neuroprogenitor stem cells
IFN: interferon
ISRE: interferon-sensitive response element
LSVs: local splicing variations
LTP: long-term potentiation
MCMV: mouse cytomegalovirus
MIA: maternal immune activation
NPCs: neural progenitor cells
NSCs: neural stem cells
ReoV: reovirus
SNHL: sensorineural hearing loss
SREs: splicing regulatory elements
UTR: untranslated region
ZIKV: Zika virus

## Declarations

Availability of data and materials: Supplementary Figures and Tables, with relevant input data files and R code, are available at GitHub repository at https://github.com/RNAdetective/Congenital_CMV-ZIKV_infections. The datasets used in this current study are publicly available in the NCBI SRA/BioProject repository, as BioProjects PRJNA422858, PRJEB38849, PRJNA487357, PRJNA358758 and PRJNA551246.

## Authors’ contributions

BWM and HP conceived the analyses and wrote the manuscript, BWM analyzed and interpreted the data and created visualizations, HM contributed expertise on ADAR editing consequences and assisted with manuscript writing. All authors read and approved the final manuscript.

## Acknowledgements

Not applicable.

## Supplementary files legends

Supplementary File 1 (XLSX). Lists of differentially expressed genes (DEGs) from comparisons of symptomatic and asymptomatic HCMV infections to control samples, from GEO2R analysis of BioProject dataset PRJNA422858 (https://www.ncbi.nlm.nih.gov/geo/geo2r/?acc=GSE108211). Sheets A, B and C show GEO2R lists of DEGs from symptomatic vs control, asymptomatic vs control, and symptomatic vs asymptomatic HCMV samples comparisons. Sheets D and E show results of Reactome pathways overrepresentation analyses for significant (FDR <= 0.05) DEGs from symptomatic vs control and asymptomatic vs control HCMV samples. Only pathways with entities FDR < 0.05 are shown.

Supplementary File 2 (CSV). (A) Characteristics of 149 editing sites from MCMV and control samples. List of ADAR edited sites (identified via chromosome (CHR) and position (POS)) and individual nucleotide counts from MCMV infections and control samples (PRJEB38849). (B) Characteristics of 21 significantly different editing sites between MCMV and control samples. List of ADAR edited sites (identified via chromosome (CHR) and position (POS)) and average editing rates from MCMV.

Supplementary File 3 (PNG). Box and violin plots of editing rates between control vs. MCMV infected mice samples (PRJEB38849); control samples are shown in red, MCMV shown in blue, respectively.

Supplementary File 4 (XLXS). Reactome pathway analysis of edited genes from infected and control samples. In bold are pathways overrepresented among editing targets with FDR < 0.1. (A) Sheet 4A shows overrepresented pathways among edited targets from MCMV and control samples (PRJEB38849). (B) Sheet 4B shows overrepresented pathways among edited targets from ZIKV and control samples (PRJNA487357). (C) Sheet 4C shows pathways among edited targets from ZIKV and control samples (PRJNA358758); there were no pathways overrepresented among editing targets with FDR < 0.1.

Supplementary File 5 (CSV). (A) Characteristics of 78 editing sites from ZIKV and control samples (PRJNA487357). List of ADAR edited sites (identified via chromosome (CHR) and position (POS)) and individual nucleotide counts from ZIKV infections and control samples. (B) Characteristics of 12 significantly different editing sites between ZIKV and control samples. List of ADAR edited sites (identified via chromosome (CHR) and position (POS)), and average editing rates from ZIKV infections and control samples (PRJNA487357).

Supplementary File 6 (PNG). Box and violin plots of editing rates between control vs. ZIKV infected mice samples (PRJNA487357); control samples are shown in red, ZIKV shown in blue, respectively.

Supplementary File 7 (XLSX). (A) Characteristics of 1276 editing sites from ZIKV and control samples. List of ADAR edited sites (identified via chromosome (CHR) and position (POS)) and individual nucleotide counts from ZIKV infections and control samples (PRJNA358758). (B) Characteristics of 148 significantly different editing sites between ZIKV and control samples. List of ADAR edited sites (identified via chromosome (CHR) and position (POS)) and average editing rates from ZIKV infections and control samples (PRJNA358758).

Supplementary File 8 (PNG). Box and violin plots of editing rates between control vs. ZIKV infected mice samples (PRJNA358758); control samples are shown in red, ZIKV shown in blue, respectively.

Supplementary File 9 (XLSX). Results of MAJIQ and VOILA analysis of editing sites from ZIKV and control samples (PRJNA358758). (A) Results of MAJIQ analysis of editing sites from ZIKV and control samples (PRJNA358758), with PSI/dPSI information for all LSVs identified. (B) Results of VOILA analysis of editing sites from ZIKV and control samples (PRJNA358758), with significant LSVs (|dPSI| > 0.2 and p < 0.05). (C) Information for LSVs in genes identified as Nova1 targets (Zhang et al., 2010).

Supplementary File 10 (XLSX). (A) Characteristics of 1355 editing sites from ZIKV and control samples (PRJNA551246). List of ADAR edited sites (identified via chromosome (CHR) and position (POS)) and individual nucleotide counts from ZIKV infections with Cambodian (FSS13025) and Brazilian ZIKV (PE243) strains and control samples. (B) Characteristics of 9 significantly different editing sites between ZIKV and control samples (PRJNA551246). List of ADAR edited sites (identified via chromosome (CHR) and position (POS)), and average editing rates from ZIKV infections with Cambodian (FSS13025) and Brazilian ZIKV (PE243) strains and control samples.

Supplementary File 11 (PNG). Box and violin plots of editing rates between control vs. ZIKV infected human induced pluripotent neuroprogenitor stem cells (hiNPCs) samples (PRJNA551246). Control samples are shown in red, ZIKV infections with Cambodian (FSS13025) and Brazilian ZIKV (PE243) strains are shown in green and blue, respectively.

Supplementary File 12 (XLSX). SubmiRine (Maxwell et al., 2015) results predicting differences in miRNA binding between unedited and edited transcripts. While no editing was found to alter miRNA binding in the MCMV or hiNPC ZIKV datasets, editing sites with potential links to changes in miRNA targeting were detected in 2 genes for the mouse ZIKV PRJNA487357 dataset and 26 genes for the mouse ZIKV PRJNA358758 dataset.

