## Supplementary figures and images for "Changes in ADAR RNA Editing Patterns in CMV and ZIKV Congenital Infections"

### SupplFile3

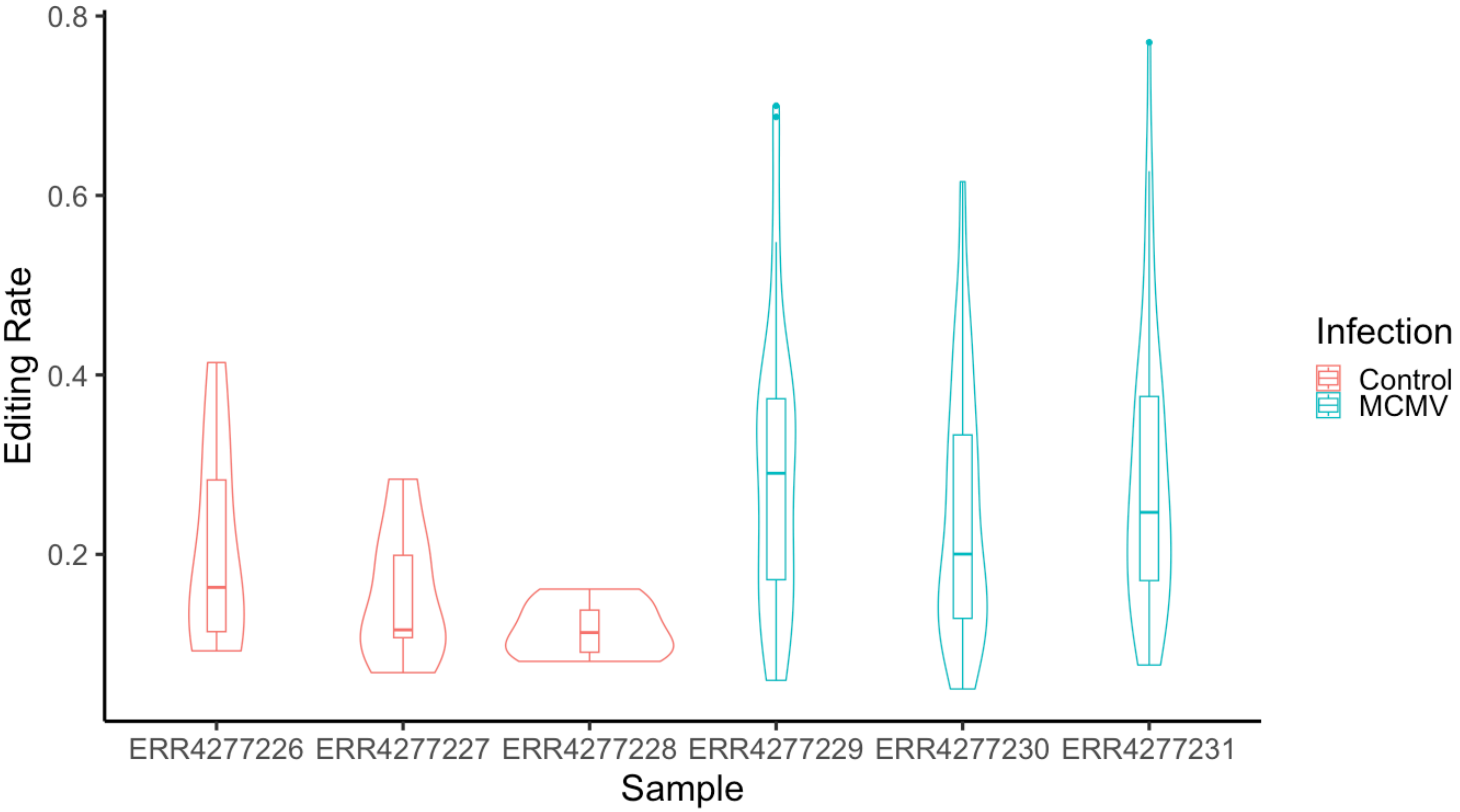

### SupplFile6

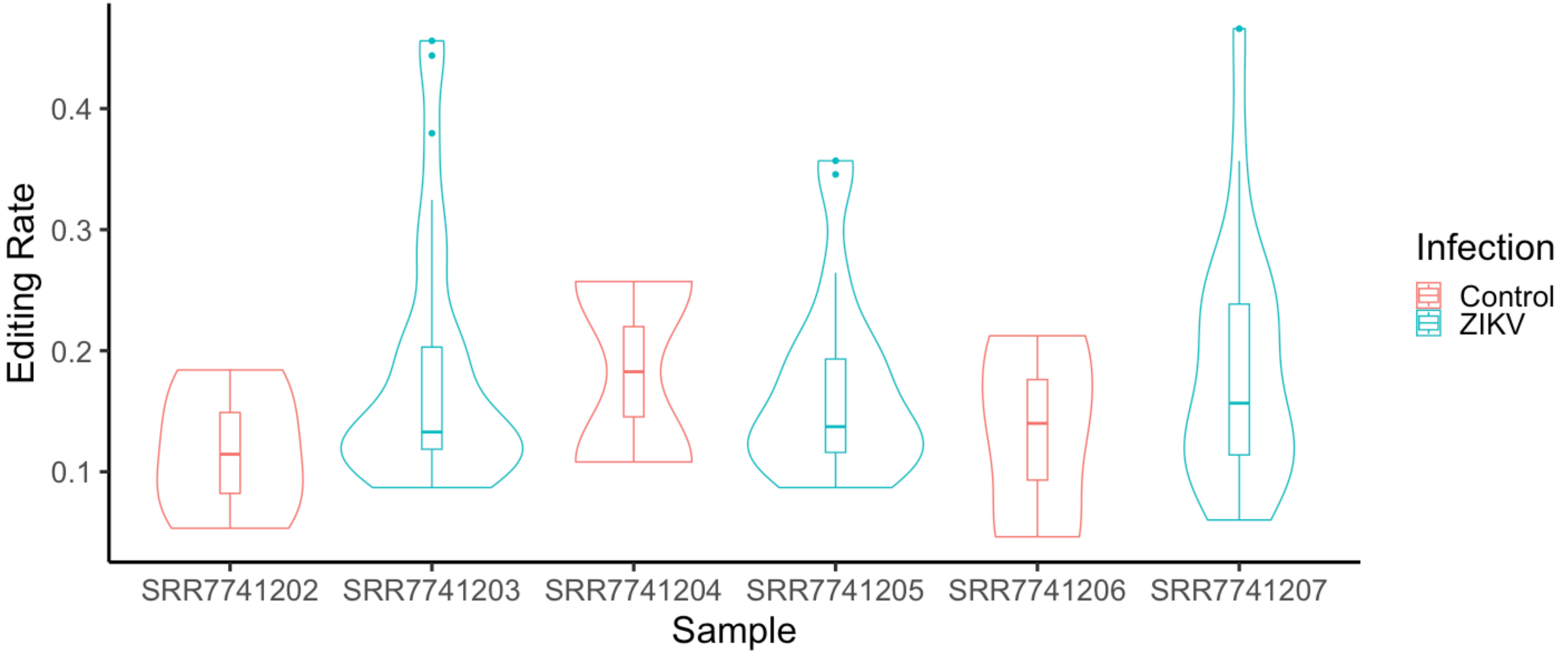

### SupplFile8

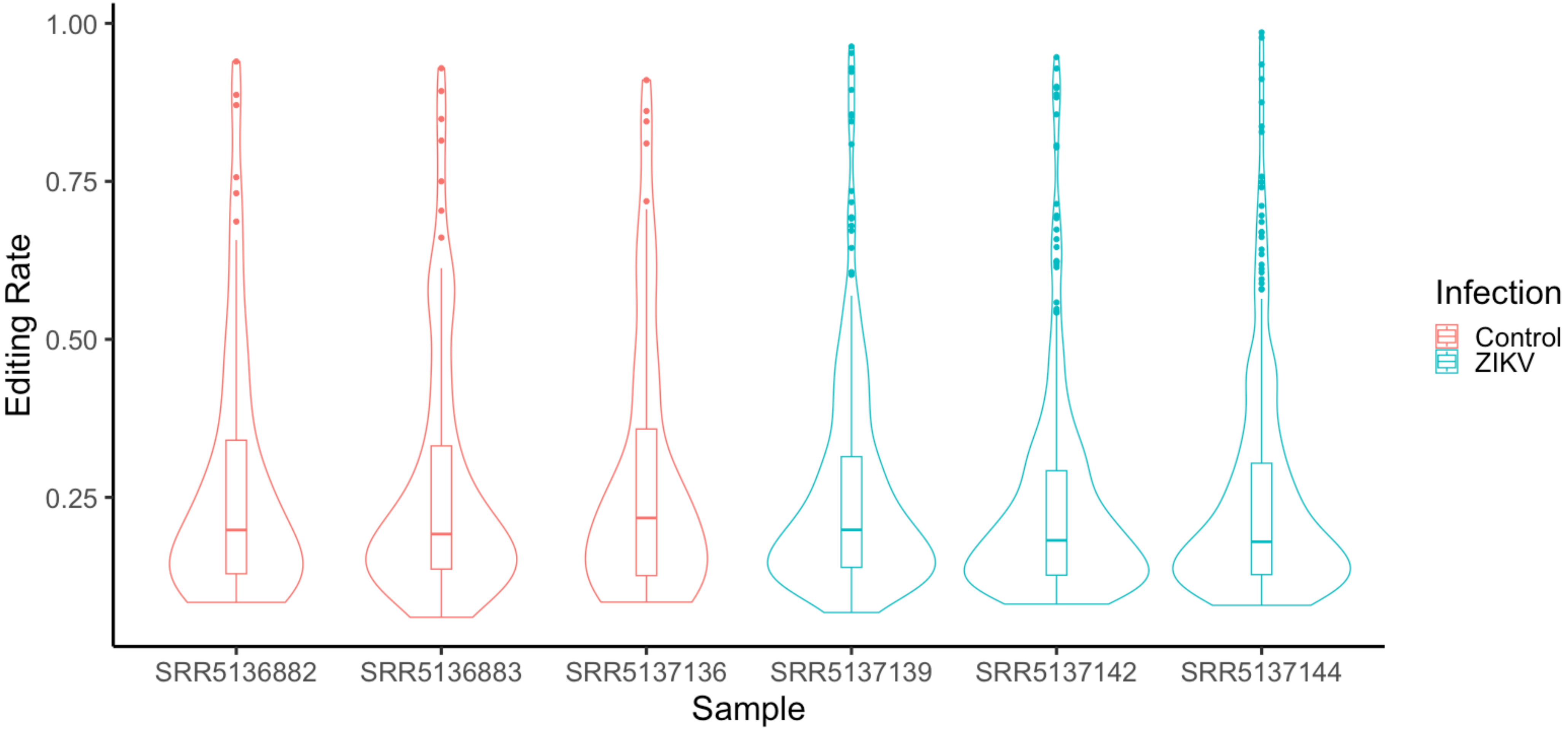

### SupplFile11

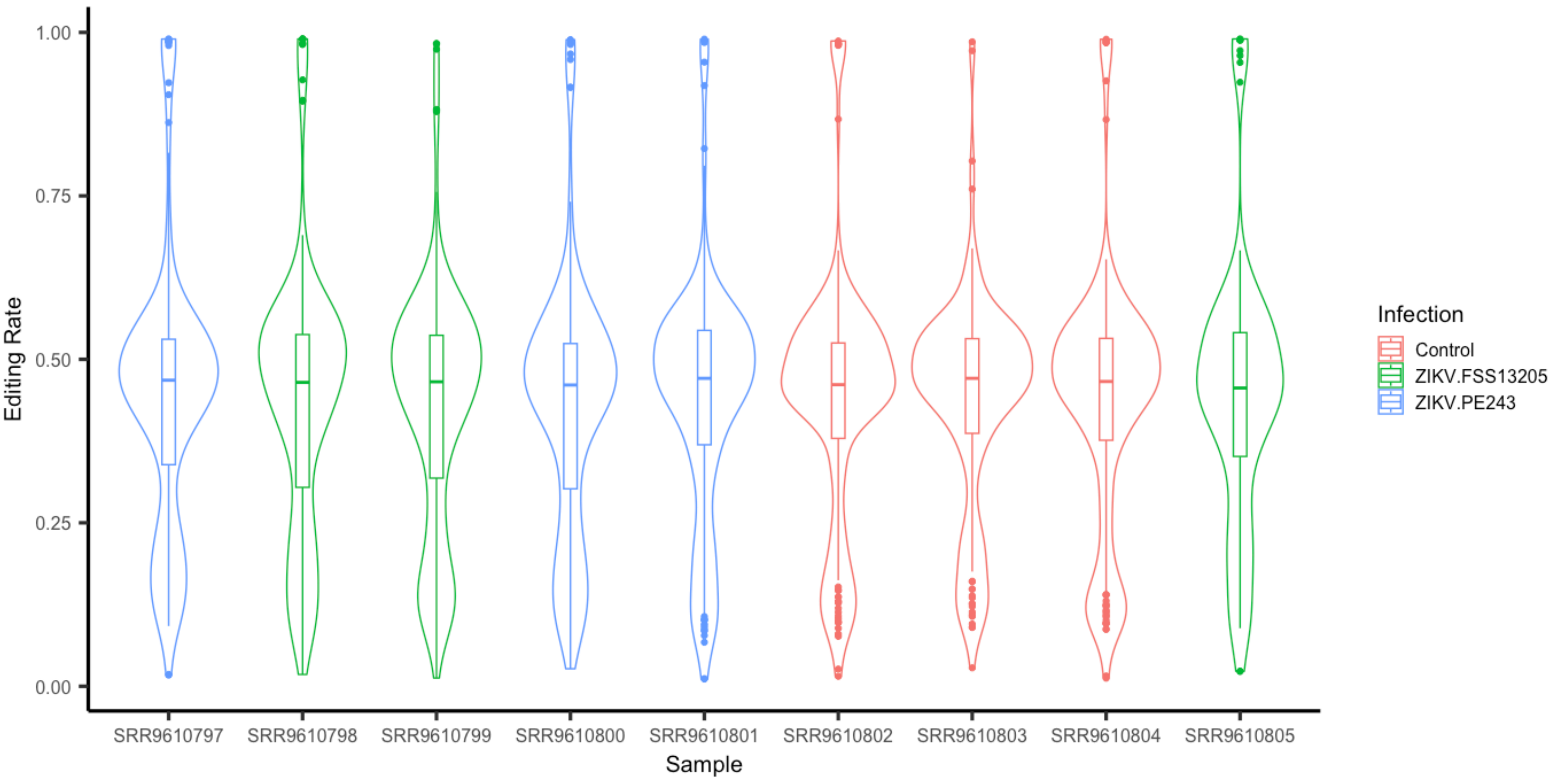
